# Heat shock protein 70 (Hsp70) is involved in the Zika virus cellular infection process

**DOI:** 10.1101/135350

**Authors:** Sujit Pujhari, Vanessa M. Macias, Ruth H. Nissly, Masashi Nomura, Suresh V. Kuchipudi, Jason L. Rasgon

## Abstract

Zika virus (ZIKV) is a historically neglected flavivirus that has recently caused epidemics in the western hemisphere. ZIKV has been associated with severe symptoms including infant microcephaly and Guillain Barré syndrome, stimulating interest in understanding factors governing ZIKV infection. Heat shock protein 70 (Hsp70) has been shown to be an infection factor for multiple viruses. We investigated the role of Hsp70 in the ZIKV infection process. We localized Hsp70 protein to the Vero cell membrane surface by confocal microscopy and demonstrated that, inside the cell, there is significant co-localization between Hsp70 and ZIKV E protein. Inducing and suppressing Hsp70 expression increased and decreased ZIKV production, respectively. Antibody blocking cell surface-localized Hsp70 decreased ZIKV cell infection rates and production of infectious virus particles, as did competition with recombinant Hsp70 protein. Our data suggest that Hsp70 is an important factor in the ZIKV infection process. Understanding the interactions between Hsp70 and ZIKV may lead to novel therapeutics for ZIKV infection, particularly for pregnant women and fetuses.

## INTRODUCTION

Zika virus (ZIKV) is a historically neglected tropical flavivirus first isolated in 1947 that typically resulted in a handful of documented cases with mild clinical phenotypes. Beginning in 2007, larger outbreaks of the virus were first recorded, culminating in a large ongoing epidemic in the western hemisphere beginning in late 2015.^1-4^ ZIKV has been declared an infection of global health concern by the World Health Organization (WHO), which is anticipating up to 4 million cases by the end of 2016.^5^ For the first time, ZIKV infection has been associated with severe symptoms including microcephaly in infants infected as fetuses, and Guillaine Barré syndrome in adults.^6,7^ The occurrence of severe clinical outcomes for fetuses and pregnant women in this outbreak has stimulated interest in determining the factors governing ZIKV infection.^8,9^

The binding of a virus to specific cell surface receptor(s) is a critical step for cellular tropism and an important determinant of pathogenesis.^10^ In general, flavivirus cell infection is mediated by an array of cell surface molecules and attachment cofactors.^11^ Heat shock protein 70 (Hsp70) has been shown to be one such factor for multiple viruses including dengue virus (DV), Japanese encephalitis virus (JEV), Hazara virus, and rotavirus, where it may act directly as a receptor or indirectly to help attach and gather viruses on the cell surface to facilitate interactions with specific high-affinity receptors.^12-15^ Recently the role of Axl, Tyro3, and TIM1 in the pathogenesis and entry of ZIKV to the neuronal and placental cell population has been described.^16-19^ However the understanding of the ZIKV cellular infection process is still in its initial stages and needs further investigation. Here we demonstrate that cell surface-bound Hsp70 is an important factor in the ZIKV cell infection process. Understanding the interactions between Hsp70 and ZIKV may lead to novel therapeutics for ZIKV infection, particularly for pregnant women and fetuses.

## RESULTS

### ZIKV binds to Hsp70 expressed on the Vero cell surface membrane

Although heat shock proteins are generally thought to be cytoplasmic, Hsp70 has been localized to the surface of the cell membranes in multiple cell types.^20^ We localized Hsp70 by IFA on the cell membrane surface of non-permeabilized Vero cells ^21^ (Figure 1). Co-staining for both Hsp70 and ZIKV at 4 hours post-infection (1 MOI) demonstrated binding of ZIKV to Hsp70 on the cell surface (Figure 1). Positive and negative staining controls are included as Figure S1.

**Figure 1.**
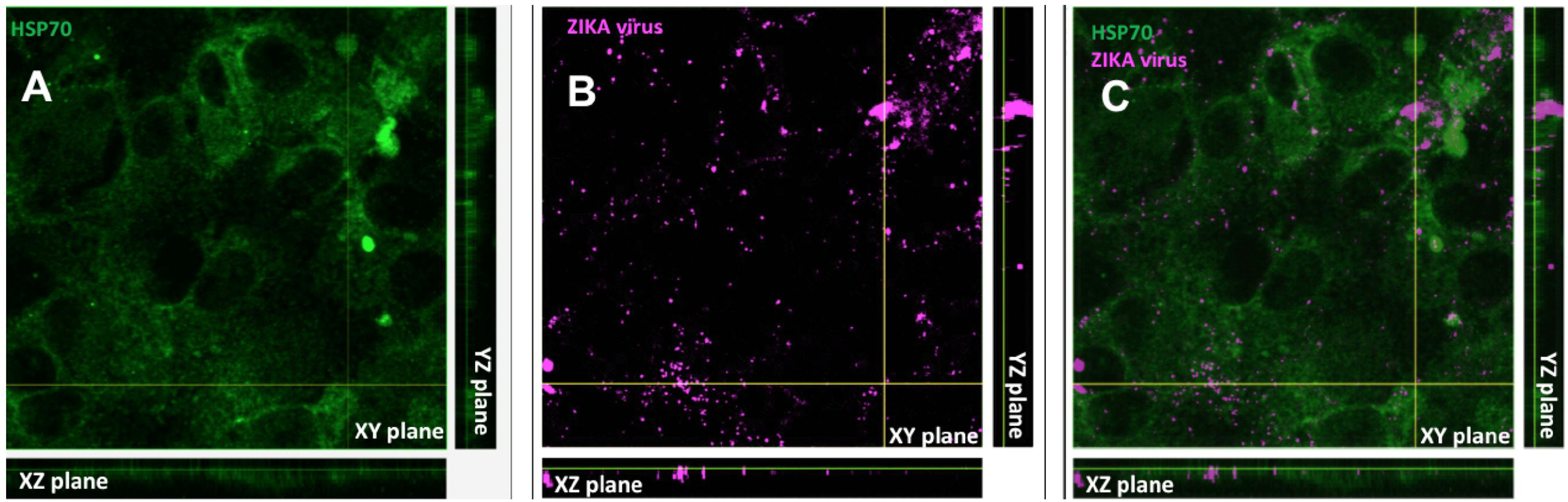
Localization of Hsp70 and ZIKV on the surface of Vero cells. A) Hsp70 (green), and B) ZIKV (purple) visualized on the surface of unpermeabilized Vero cells by confocal microscopy. (C) Merge.

### Intracellular localization of HSP70 and ZIKV E protein

The E glycoproteins are projected out of the ZIKV virion surface, making them potential targets for interaction with Hsp70. We used imaging flow cytometry and confocal microscopy to investigate this interaction. Images showed qualitatively similar distribution patterns of Hsp70 and ZIKV E protein intracellularly in infected cells by both confocal microscopy and imaging FACS analysis (Figure 2). Confocal imaging and co-localization analysis of intracellularly stained cells demonstrated a high degree of correlation between ZIKV E protein and Hsp70 (correlation coefficient = 0.91 ± 0.04, compared to 0.38 ± 0.09 for randomly selected areas of the images) (Figure 2C).

**Figure 2.**
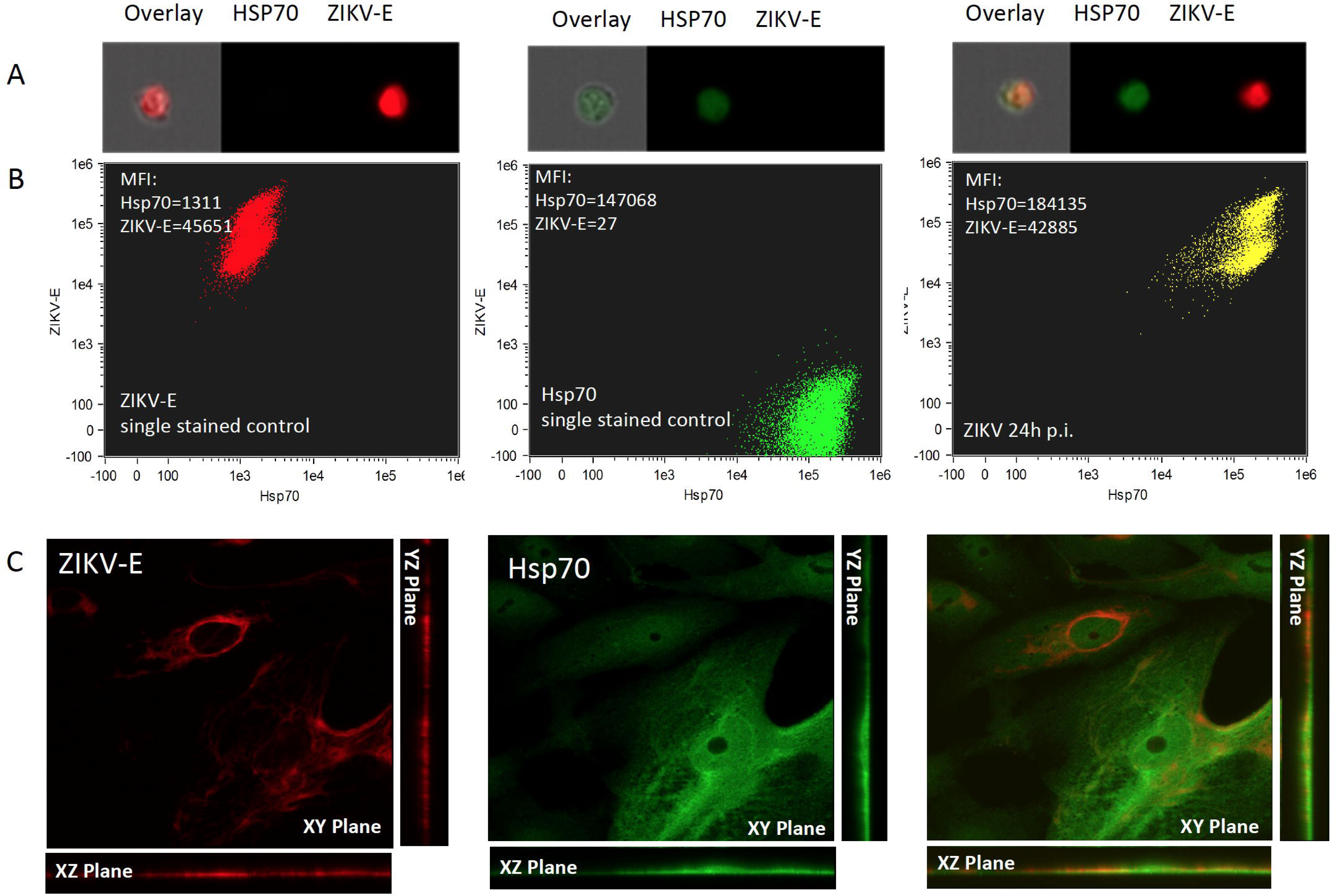
ZIKV-E protein interacts with Hsp70. A) Single-cell imaging flow cytometry images of permeabilized cells demonstrating intracellular colocalization of Hsp70 (green; Alexa Fluor 488) and ZIKV E protein (red; Alexa Fluor 633). B) Flow cytometry analysis of permeabilized cells stained for Hsp70 (green) and ZIKV-E protein (red). C) Confocal microscopy images of permeabilized cells demonstrating co-localization of intracellular Hsp70 (green) and ZIKV E protein (red).

### Hsp70 protein expression modulates ZIKV infection

To investigate the role of Hsp70 in ZIKV infection, we induced Hsp70 in Vero cells by both elevated temperature and chemical induction, infected cells with ZIKV, and measured the production of infectious ZIKV in the culture supernatant by plaque assay. Upon heat treatment, Hsp70 protein levels were induced approximately 2-fold at 8 hours post heat treatment compared with non-heat treated cells (Figure 3A). For chemical induction, we treated cells with geldanamycin (which inhibits Hsp90 and stabilizes Hsp70 mRNA leading to increased expression of Hsp70 protein).^22^ Hsp70 protein expression was induced approximately 7-fold in a dose-dependent manner 24 hours post geldanamycin treatment compared with untreated cells (Figure 3B). In both heat-treated and chemically induced cells, infectious ZIKV was increased in the supernatant compared to non-induced cells (Figure 3C-D). We then inhibited Hsp70 by treating cells with the inhibitor rhodocyanine MKT-077, which is an allosteric inhibitor of Hsp70 that preferentially binds and inhibits the ADP-bound forms of Hsp70.^23^ Production of infectious ZIKV was reduced in treated cells compared to controls (Figure 3E). Minimal dose-dependent geldanamycin and MKT-077 cell toxicity were observed in cells at the concentration ranges used in these experiments (Figure S2) and is similar to that seen in other studies.^24-26^

**Figure 3.**
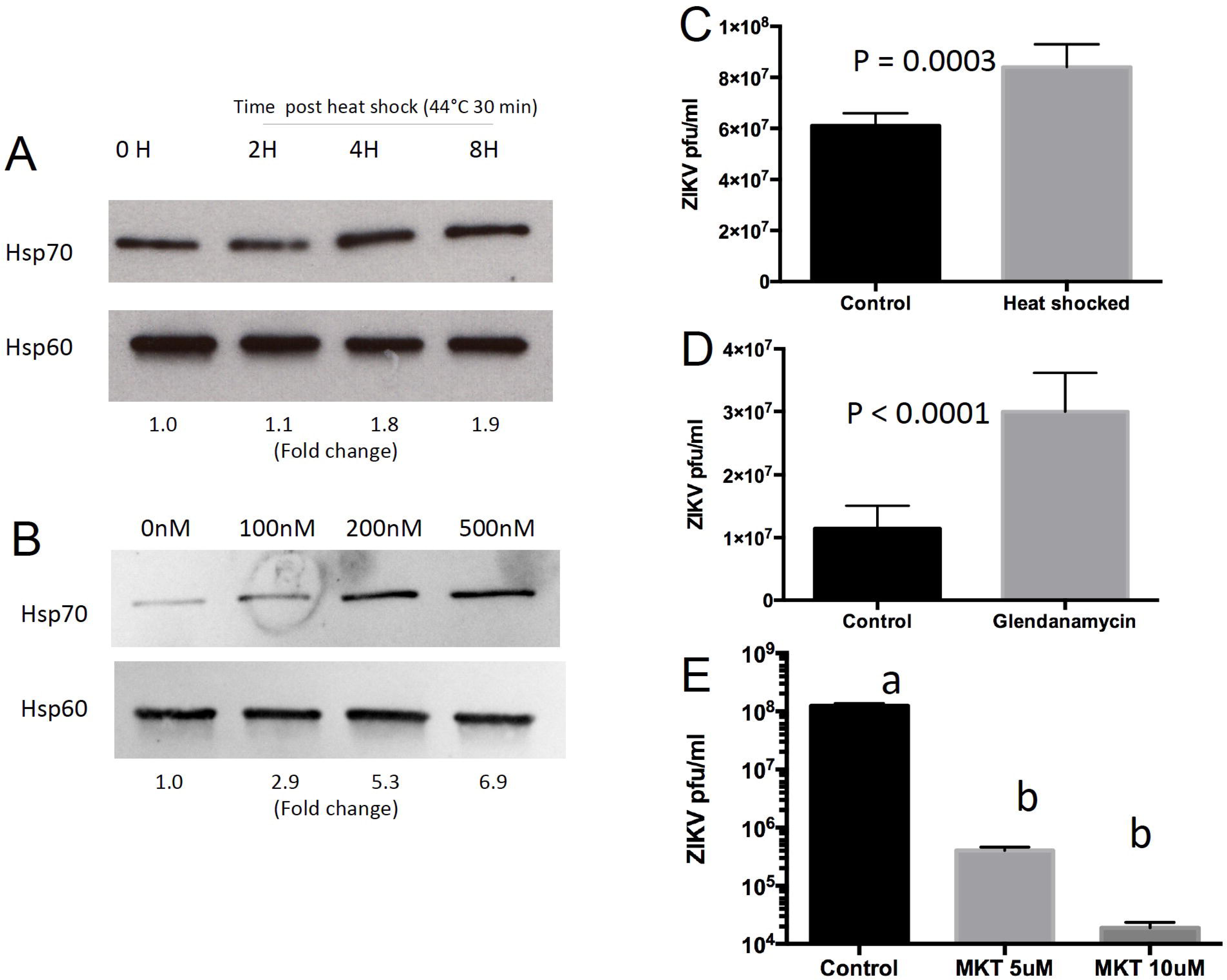
Heat shock 70 (Hsp70) protein facilitates ZIKV infection. Heat (A) and geldanamycin (B) induction of Hsp70 protein in Vero cells. Induction fold-change was calculated by quantitating normalized band intensities using ImageJ software. ZIKV is induced by heat (C) and geldanamycin treatment (D) in Vero cells. (E) Treatment of Vero cells with the Hsp70 inhibitor MKT077 suppresses ZIKV. Treatments with different lowercase letters are significantly different at P < 0.0001. Error bars denote 95% confidence intervals. Uncropped Western blots scanned from notebook are available as Figure S3. MKT = MKT077.

### Antibody blocking of cell surface Hsp70 reduces ZIKV infection

Based on these results, we hypothesized that surface-localized Hsp70 might be involved in the ZIKV entry process. Vero cells were blocked with antibodies to Hsp70, β-actin (negative control) or Axl (which has been previously demonstrated to act as a cell receptor for ZIKV ^17^ and which served as a positive control for the assay). Blocking with anti-Hsp70 antibody reduced the number of ZIKV infected cells (Figure 4A) and reduced the number of plaques by plaque reduction neutralization assay to a similar extent as blocking Axl (Figure 4B).

**Figure 4.**
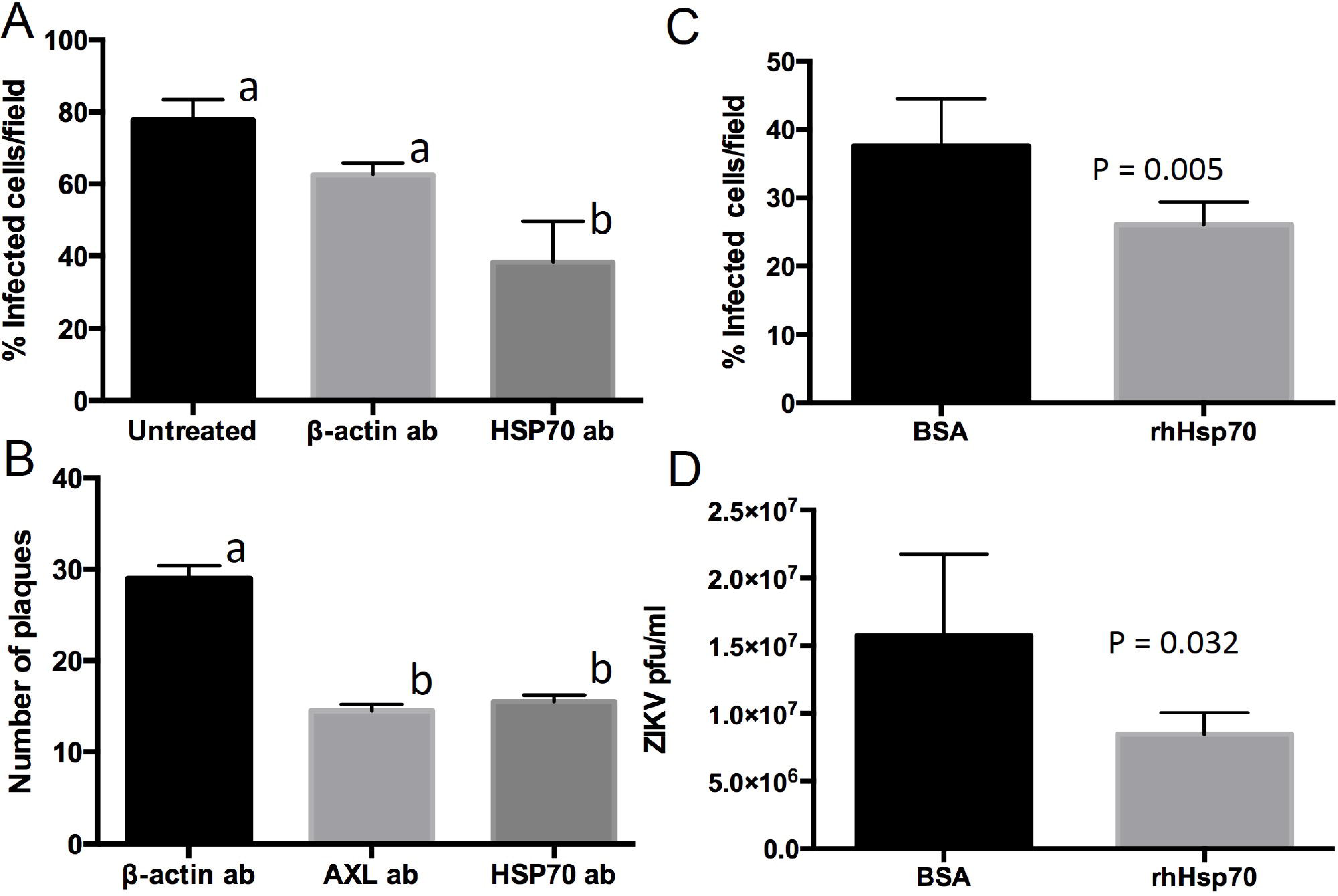
Blocking ZIKV cellular attachment to Hsp70 reduces cell infection rates and infectious particle production. Blocking Vero cells with Hsp70 antibody reduces percentage of ZIKV-infected cells (A) and numbers of plaques by plaque reduction neutralization assay (B). Competitive blocking with recombinant human Hsp70 protein (rhHsp70) reduces both the percentage of infected cells (C) and production of infectious ZIKV particles by plaque assay (D). Error bars denote 95% confidence intervals. Treatments with different lowercase letters are statistically different at P < 0.01.

### Competition for ZIKV attachment with recombinant Hsp70 protein reduces ZIKV infection

We treated cells with recombinant human Hsp70 (rhHsp70) as competitor for the putative cell attachment protein(s) present on the ZIKV infectious particle. ZIKV was incubated with rhHsp70 and used to infect Vero cells, after which ZIKV production was assayed by plaque assay and by quantifying the number of infected cells by immunofluorescence microscopy. Incubating ZIKV with rhHsp70 reduced the number of infected cells post-infection (Figure 4C) and reduced the amount of infectious virus in the supernatant compared to BSA-treated controls (Figure 4D).

## DISCUSSION

Viruses, as obligate intra-cellular parasites, entirely depend on host cellular factors for cell entry, genome replication, protein synthesis, and virion assembly, and have evolved strategies to counter cellular defense mechanisms to initiate and establish infection. Hsps have been found to be critical factors for the pathogenesis of many viruses.^27^ For example, HIV, rotavirus, and influenza virus replication are negatively regulated by Hsp70, whereas dengue, Japanese encephalitis, hepatitis C and Hazara viruses utilize Hsp70 for replication.^12, 15, 28-33^

Until recently, ZIKV was not considered a serious public health threat, and thus its biology has not been studied in great detail. Previous work identified Axl, Tyro3, and TIM1 as major cellular players in the ZIKV cell entry process. However, Cas9-mediated deletion of Axl-1 in human neuronal progenitor cells and cerebral organoids did not protect these cells from ZIKV infection.^34^ This suggests that ZIKV uses multiple classes of cell surface molecules as receptors in different cell types. As an initial event in the infection process, virus particles attach and aggregate on the cell surface, providing a window of opportunity for the virus to bind to a specific receptor(s) on the dynamic cell membrane. Other molecules (including Hsp70) have been implicated for the attachment and aggregation of other viruses.^11, 24, 35^ Although Hsp70 chaperones do not contain export signal peptide sequences, they are found on the cell surface in clathrin coated pits and within endosome/lysosome-related vesicles.^36^ They translocate spontaneously from the cytosol into the plasma membrane after oligomerization and binding to phosphatidylserine.^20^

We detected the presence of Hsp70 on the cell surface of Vero cells and observed co-localization between ZIKV and Hsp70 on the surface of the cell membrane (Figure 1). In the cases of JEV and DV infection, surface-localized Hsp70 interacts with the viral E protein and acts as a putative receptor.^13, 14^ In the early stage of JEV infection, the E protein interacts with lipid rafts and also activates the phosphoinositide 3-kinase/Akt signaling pathway to promote the cell entry event.^37^ In addition, cytoplasmic Hsp70 has been shown to modulate the replication cycle of a related Flavivirus (HCV) at the translation and virion assembly stages^38^. Our observations of an interaction between Hsp70 and ZIKV E protein (Figure 2) are indicative of similar processes during ZIKV infection.

Hsp70 expression increases after ZIKV infection (Figure 3A) similar to JEV, West Nile virus (WNV) and DV. Since a large number of viral proteins are synthesized in a relatively short period of time, the virus may need chaperone molecules to assist and facilitate the protein folding process. In this study we found that inducing Hsp70 either physically or chemically resulted in increased production of ZIKV in infected cells, while chemical inhibition of Hsp70 with MKT077 showed the opposite effect (Figure 3). A possible explanation of the decrease in virus titer with suppression of Hsp70 could be the decrease in the stability of the ZIKV proteins in the absence of the chaperone molecule (Hsp70) that results in activation of proteasomal degradation, similar to results seen with Hepatitis C virus (HCV) and JEV infections.^25, 33^

Based on these results, we designed a receptor blocking experiment to ascertain the role of Hsp70 in the cellular entry process of ZIKV. Antibody blocking of cell surface-localized Hsp70 or pre-incubation of ZIKV with recombinant Hsp70 significantly reduced ZIKV infection (Figure 4). Specifically, reduction in the number of infected cells and reduction in viral plaque forming units suggests that initial viral infection was inhibited, consistent with the hypothesis that Hsp70 is acting as a cell surface receptor for ZIKV attachment and invasion.

Due to the involvement of Hsp70 in the infection processes of multiple viruses, it has been suggested that this protein may be a potential molecular target for anti-viral therapies.^39-41^ In particular, the association between Hsp70 and ZIKV suggests a critical role for this interaction during the viral infection and/or replication process. Our data suggest that Hsp70 is one of several critical factors mediating the initial entry of ZIKV into host cells. As there is currently no effective therapy or vaccine for ZIKV infection, novel therapies to reduce infection and severe clinical outcomes, particularly for pregnant women and developing fetuses, are of extreme importance. The results of this work validate Hsp70 as a potential target for future anti-ZIKV therapies, if detrimental host effects of Hsp70 blockage can be overcome.

## MATERIALS AND METHODS

Cells and virus: Vero cells were obtained from ATCC and were grown in complete Dulbecco’s modified Eagle’s medium (DMEM) supplemented with 10% fetal bovine serum (FBS; Invitrogen), 1X penicillin and streptomycin (Invitrogen) and 2 mM Lglutamine (Invitrogen). The MR766 ZIKV strain was obtained from BEI resources, propagated in Vero cells, and titers determined by standard plaque assays.

Antibodies and chemical reagents: The E glycoprotein of ZIKV was detected using anti-flavivirus group antigen antibody, clone D1-4G2-4-15 (4G2, Millipore). Rabbit polyclonal Hsp70 antibody was purchased from Abcam (ab79852). Rabbit Hsp60 antibody (SAB4501464) was purchased from Sigma. Alexa 488-conjugated anti-mouse and anti-rabbit antibodies (A11001 and A11034) were purchased from Life Technologies. Goat β-actin antibody (ab8229) was purchased from Abcam. Rabbit Axl antibody (C89E7) was purchased from Cell Signaling. MKT-077 and Geldanamycin were purchased from Sigma.

Immuno-localization of Hsp70 on and in Vero cells: To analyze co-expression of Hsp70 and ZIKV E glycoprotein by flow cytometry, 6x10^5^ Vero cells were plated and cultured overnight on 6-well plates. After being washed twice with culture medium without fetal bovine serum (FBS), cell monolayers were infected with 0.1 ml of medium containing 5 MOI of ZIKV. Mock infection controls were performed without the virus. After 90 minutes of adsorption, unabsorbed virus particles were removed and 2 ml of maintenance medium added. Virus and mock-infected cells were harvested (using 0.25% trypsin) at 24h and 48h post-infection for flow cytometry analysis. The staining process was performed at 4°C using ice-cold reagents. Cells were washed twice (500 × g, 5 minutes) with medium containing 10% FBS to neutralize trypsin, then with flow buffer (PBS with 0.1% BSA). Cells were fixed (4% freshly prepared paraformaldehyde in PBS), suspended in permeabilization buffer (0.5% saponin in flow buffer), for 10 minutes, and incubated with Hsp70 and Zika virus E protein primary antibodies diluted 1:100 in permeabilization buffer for 60 minutes. Cells were then washed twice with permeabilization buffer and incubated for 30 minutes with anti-rabbit-Alexa Fluor 488 and anti-mouse-Alexa Fluor 633 conjugated secondary antibodies diluted 1:200 in permeabilization buffer. Cells were washed twice with flow buffer, and resuspended in 2% paraformaldehyde and stored at 4°C until analysis. Cells were analyzed using an Amnis FlowSight (EMD Millipore, USA) with 488 nm, 633 nm, and 785 nm laser excitation. 20,000 images of non-debris objects were collected for each sample using INSPIRE® software. In-focus cells from data files were analyzed using IDEAS® version 6.2 (Amnis Corporation, USA) by gating on non-autofluorescent single cells.

Immunofluorescence microscopy was performed on Vero cells to elucidate the distribution of Hsp70. Cells were seeded at a density of 2 × 10^5^ cells/well in a 2 well chamber slide at 37 °C. After 24 h, cells were fixed with 4% paraformaldehyde at room temperature. Cells were permeabilized to investigate Hsp70/ZIKV co-localization inside the cell or not permeabilized to investigate Hsp70 distribution on the cell surface. Hsp70 was stained using anti-Hsp70 antibody and examined by confocal microscopy.

Plaque assays: Vero cells (5×10^5^ cells/well) were grown to a confluent monolayer in 6 well plates and infected with serial dilutions of ZIKV-infected culture supernatant. Incubation was carried out for 2 hours at 37 °C, after which monolayers were rinsed with sterile PBS. Monolayers were overlaid with maintenance medium containing 0.6% molten agarose and incubated at 37 °C for 3 days. At the end of incubation period, secondary overlay media (primary overlay media along with 1% neutral red) was poured and incubated at 37 °C overnight. The next day plaques were counted.

Plaque reduction neutralization assays: Confluent cell monolayers were grown in 6 well plates and incubated with 1:50 dilution of polyclonal β-actin, Axl or Hsp70 antibodies at 4°C for 1h, rinsed with sterile PBS, and incubated with 10^5^ pfu ZIKV particles, and processed as described above for plaque assays.

Western blots: Cells were washed twice with PBS and lysed in RIPA buffer (Sigma). The lysate was cleared by centrifugation at 14,000 rpm for 20 min at 4 °C and subjected to 10% sodium dodecyl sulphate polyacrylamide gel electrophoresis (SDS/PAGE), after which proteins were transferred to nitrocellulose membranes (BioRad) that were blocked for 60 min at room temperature in 1X TBST buffer (50 mM Tris-HCl pH7.4, 250 mM NaCl, 0.1% Tween-20) containing 5% non-fat milk powder. Membranes were incubated overnight at 4 °C with anti-Hsp60 or anti-Hsp70 primary antibodies (1:1000). The next day, blots were washed 5 times with 1X TBST, and incubated with the corresponding detection antibody (1:2000) in 1X TBST containing 1% milk solution at room temperature for 1 h. Signals were detected with the enhanced chemiluminescence method (GE healthcare) or AP-based colorimetric kit (BioRad).

For quantitation, blots were scanned and band intensities quantified using ImageJ software (NIH, Bethesda, MD, USA). The background was first subtracted from the gel image, the intensity of the Hsp70 band and Hsp60 control band quantified for each sample by measuring the area under the intensity curve, and the value of Hsp70 divided by the value for Hsp60. Quotients for each sample were normalized to hour 0 (for heat treatment) or untreated (for geldanamycin) values to calculate protein induction. Uncropped blots are included as Figure S3.

Cell heat shock assay: Confluent cell monolayers were grown in 12 well plates and incubated at 44°C for 20 min or at 37°C (negative control). After heat shock, cells were allowed to recover for 30 min at 37°C. Cells were harvested 2h, 4h and 8h post heat treatment to check the expression level of Hsp70. At 8h post heat shock cells were infected with ZIKV at an MOI of 0.1. Culture supernatants were harvested 48h post infection and titrated for infectious ZIKV by plaque assay.

MTT assay: To determine cell cytotoxicity, the colorimetric MTT assay (Biovision Inc, USA) was used. Metabolically active, viable cells convert MTT into formazan, a water insoluble product; however, dead cells lose this ability. Vero cells (1 × 10^4^ cells/well) were cultured in a 96-well plate at 37 °C, and exposed to varying concentrations of geldanamycin and MKT-077 for 24 h. Cells treated with medium only served as the control group. After removing the supernatant of each well and washing twice by PBS, 20 μl of MTT reagent (5 mg/ml) and 100 μl of medium were introduced. Following incubation for another 3.5 h, the resultant formazan crystals were dissolved in MTT solvent (100 μl) and the absorbance intensity measured by a microplate reader at 600 nm. All experiments were performed in triplicates, and relative cell viability (%) was expressed as a percentage relative to the untreated control cells.

Chemical Hsp70 induction assay: Cells were grown in 96 well plates and treated with 100nM, 200nM, or 500nM geldanamycin in DMEM for 1 hour at 37 °C. Cells treated with DMEM lacking geldanamycin served as a negative control. After treatment, cells were washed twice with 1X PBS and infected with ZIKV at an MOI of 0.1. After 1 hour at 37 °C in 5% CO 2, the virus inoculum was removed and replaced with 100 μl of growth medium. Culture supernatants were harvested 48h post infection and titrated by plaque assay.

MKT077 Hsp70 inhibition assay: Cells were grown in 96 well plates and treated with 5μM or 10μM MKT077in DMEM for 1 hour at 37°C. Cells treated with DMEM with no MKT700 served as a negative control. After treatment, virus infection and quantitation were carried out as described above.

Antibody blocking Hsp70 on the cell surface and ZIKV infection assays: Vero cells were cultured in Permanox two-well chamber slides (Thermo Fisher Scientific) at a density of 2x10^5^ and incubated with a 1:50 dilution of Hsp70 polyclonal antibody at 4°C for 2 h. β-actin antibody (1:50) was used as a negative control, and untreated cells were included as an additional negative control. After 2 hours, cells were washed with phosphate buffered saline (PBS) and infected with ZIKV at an MOI of 2.5 and incubated at 37°C in 5% CO_2_. After 24 h, cells were fixed with 4% formaldehyde in PBS and permeabilized with 0.1% triton X-100 in PBS for 10 min at room temperature. Prior to staining, cells were blocked with 5% BSA to prevent non-specific binding of antibodies and conjugate. Cells were incubated with primary antibody targeting the flavivirus E protein at 1:100 dilution for 2 h at room temperature, followed by washing with PBS and incubation with Alexa Fluor-488 conjugated antibodies at a 1:500 dilution for 1 h at room temperature. Cells were incubated with 0.5 μg/ml of 4′,6-diamidino-2-phenylindole (DAPI; Life Technologies) at 37°C for 10 min to stain cell nuclei. Slides were washed once with PBS, once with deionized water, and then were air-dried and mounted using ProLong Gold Antifade Reagent (Life Technologies) prior to examination on a fluorescence microscope (Olympus BX41). For each slide, 5-10 random fields were chosen and for each field the number of infected cells (counted by DAPI-stained nuclei) were counted, as well as the number of cells infected with ZIKV to get the percentage of infected cells per field.

In a separate experiment, cells were blocked with β-actin antibody (negative control) or with Axl antibody (positive control), or Hsp70 polyclonal antibody and cells infected and incubated as described above. The culture supernatant was removed from cells and infectious ZIKV particles in each treatment quantified by plaque reduction neutralization assays as described above.

Competition assay: To block virus surface ligands, 10^5^ PFU of ZIKV was incubated with 1000 ng recombinant human Hsp70 (rhHsp70) protein (Enzo Life Science) or BSA (control) for 90 minutes on ice. Vero cells were infected with ZIKV preparations at an MOI of 1 at 37°C for 1 h, after which cells were washed three times with culture medium. Fresh medium was added to infected cells which were then incubated at 48 h at 37°C. The percentage of infected cells was determined by immuno-fluorescent microscopy, and infectious ZIKV particles in the culture supernatant were determined by plaque assays as described above.

Statistical analysis: Experiments with paired treatments were analyzed by t-tests. Experiments with 3 treatments were analyzed by analysis of variance with Bonferroni’s correction for multiple tests. Experiments were replicated two to three times.

## ACKNOWLEDGEMENTS

This study was supported by funds from the Penn State Huck Institutes of the Life Sciences, and NIH/NIAID grants R21AI128918 and R01AI116636 to JLR. The authors wish to thank the staff of the Flow Cytometry Lab of the Huck Institutes Microscopy and Cytometry Facility for their assistance with flow cytometry experiments. We are also thankful to Missy L Hazen for her technical support in confocal imaging.

## REFERENCES CITED

1. Lanciotti RS, Kosoy OL, Laven JJ, Velez JO, Lambert AJ, Johnson AJ, et al. Genetic and serologic properties of Zika virus associated with an epidemic, Yap State, Micronesia, 2007. Emerg Infect Dis 2008; 14:1232–9.

2. Hayes EB. Zika virus outside Africa. Emerg Infect Dis 2009; 15:1347–50.

3. Dyer O. Zika virus spreads across Americas as concerns mount over birth defects. BMJ 2015; 351:h6983.

4. Duffy MR, Chen TH, Hancock WT, Powers AM, Kool JL, Lanciotti RS, et al. Zika virus outbreak on Yap Island, Federated States of Micronesia. N Engl J Med 2009; 360:2536–43.

5. Gulland A. Zika virus is a global public health emergency, declares WHO. BMJ 2016; 352:i657.

6. Carteaux G, Maquart M, Bedet A, Contou D, Brugieres P, Fourati S, et al. Zika Virus Associated with Meningoencephalitis. N Engl J Med 2016; 374:1595–6.

7. de Paula Freitas B, de Oliveira Dias JR, Prazeres J, Sacramento GA, Ko AI, Maia M, et al. Ocular Findings in Infants With Microcephaly Associated With Presumed Zika Virus Congenital Infection in Salvador, Brazil. JAMA Ophthalmol 2016.

8. Brasil P, Pereira JPJr., Raja Gabaglia C, Damasceno L, Wakimoto M, RM Ribeiro Nogueira, et al. Zika Virus Infection in Pregnant Women in Rio de Janeiro -Preliminary Report. N Engl J Med 2016.

9. Cao-Lormeau VM, Blake A, Mons S, Lastere S, Roche C, Vanhomwegen J, et al. Guillain-Barre Syndrome outbreak associated with Zika virus infection in French Polynesia: a case-control study. Lancet 2016; 387:1531–9.

10. Schneider-Schaulies J. Cellular receptors for viruses: links to tropism and pathogenesis. J Gen Virol 2000; 81:1413–29.

11. Cruz-Oliveira C, Freire JM, Conceicao TM, Higa LM, Castanho MA, AT Da Poian. Receptors and routes of dengue virus entry into the host cells. FEMS Microbiol Rev 2015; 39:155–70.

12. Broquet AH, Lenoir C, Gardet A, Sapin C, Chwetzoff S, Jouniaux AM, et al. Hsp70 negatively controls rotavirus protein bioavailability in caco-2 cells infected by the rotavirus RF strain. J Virol 2007; 81:1297–304.

13. Das S, Laxminarayana SV, Chandra N, Ravi V, Desai A. Heat shock protein 70 on Neuro2a cells is a putative receptor for Japanese encephalitis virus. Virology 2009; 385:47–57.

14. Reyes-Del Valle J, Chavez-Salinas S, Medina F, RM Del Angel. Heat shock protein 90 and heat shock protein 70 are components of dengue virus receptor complex in human cells. J Virol 2005; 79:4557–67.

15. Surtees R, Dowall SD, Shaw A, Armstrong S, Hewson R, Carroll MW, et al. Heat Shock Protein 70 Family Members Interact with Crimean-Congo Hemorrhagic Fever Virus and Hazara Virus Nucleocapsid Proteins and Perform a Functional Role in the Nairovirus Replication Cycle. J Virol 2016; 90:9305–16.

16. Mlakar J, Korva M, Tul N, Popovic M, Poljsak-Prijatelj M, Mraz J, et al. Zika Virus Associated with Microcephaly. N Engl J Med 2016; 374:951–8.

17. Nowakowski TJ, Pollen AA, Di Lullo E, Sandoval-Espinosa C, Bershteyn M, Kriegstein AR. Expression Analysis Highlights AXL as a Candidate Zika Virus Entry Receptor in Neural Stem Cells. Cell Stem Cell 2016; 18:591–6.

18. Savidis G, McDougall WM, Meraner P, Perreira JM, Portmann JM, Trincucci G, et al. Identification of Zika Virus and Dengue Virus Dependency Factors using Functional Genomics. Cell Rep 2016; 16:232–46.

19. Tabata T, Petitt M, Puerta-Guardo H, Michlmayr D, Wang C, Fang-Hoover J, et al. Zika Virus Targets Different Primary Human Placental Cells, Suggesting Two Routes for Vertical Transmission. Cell Host Microbe 2016.

20. Vega VL, Rodriguez-Silva M, Frey T, Gehrmann M, Diaz JC, Steinem C, et al. Hsp70 translocates into the plasma membrane after stress and is released into the extracellular environment in a membrane-associated form that activates macrophages. J Immunol 2008; 180:4299–307.

21. Hinners I, Moschner J, Nolte N, Hille-Rehfeld A. The orientation of membrane proteins determined in situ by immunofluorescence staining. Anal Biochem 1999; 276:1–7.

22. Elo MA, Kaarniranta K, Helminen HJ, Lammi MJ. Hsp90 inhibitor geldanamycin increases hsp70 mRNA stabilisation but fails to activate HSF1 in cells exposed to hydrostatic pressure. Biochim Biophys Acta 2005; 1743:115–9.

23. Miyata Y, Li X, Lee HF, Jinwal UK, Srinivasan SR, Seguin SP, et al. Synthesis and initial evaluation of YM-08, a blood-brain barrier permeable derivative of the heat shock protein 70 (Hsp70) inhibitor MKT-077, which reduces tau levels. ACS Chem Neurosci; 4:930–9.

24. Li YH, Tao PZ, Liu YZ, Jiang JD. Geldanamycin, a ligand of heat shock protein 90, inhibits the replication of herpes simplex virus type 1 in vitro. Antimicrob Agents Chemother 2004; 48:867–72.

25. Li YH, Lu QN, Wang HQ, Tao PZ, Jiang JD. Geldanamycin, a ligand of heat shock protein 90, inhibits herpes simplex virus type 2 replication both in vitro and in vivo. J Antibiot (Tokyo) 2012;65:509–12.

26. Taguwa S, Maringer K, Li X, Bernal-Rubio D, Rauch JN, Gestwicki JE, et al. Defining Hsp70 Subnetworks in Dengue Virus Replication Reveals Key Vulnerability in Flavivirus Infection. Cell 2015; 163:1108–23.

27. Mayer MP. Recruitment of Hsp70 chaperones: a crucial part of viral survival strategies. Rev Physiol Biochem Pharmacol 2005; 153:1–46.

28. Chen YJ, Chen YH, Chow LP, Tsai YH, Chen PH, Huang CY, et al. Heat shock protein 72 is associated with the hepatitis C virus replicase complex and enhances viral RNA replication. J Biol Chem 2010; 285:28183–90.

29. Hirayama E, Atagi H, Hiraki A, Kim J. Heat shock protein 70 is related to thermal inhibition of nuclear export of the influenza virus ribonucleoprotein complex. J Virol 2004; 78:1263–70.

30. Lordanskiy S, Zhao Y, Dubrovsky L, Iordanskaya T, Chen M, Liang D, et al. Heat shock protein 70 protects cells from cell cycle arrest and apoptosis induced by human immunodeficiency virus type 1 viral protein R. J Virol 2004; 78:9697–704.

31. Li G, Zhang J, Tong X, Liu W, Ye X. Heat shock protein 70 inhibits the activity of Influenza A virus ribonucleoprotein and blocks the replication of virus in vitro and in vivo. PLoS One 2011; 6:e16546.

32. Padwad YS, Mishra KP, Jain M, Chanda S, Ganju L. Dengue virus infection activates cellular chaperone Hsp70 in THP-1 cells: downregulation of Hsp70 by siRNA revealed decreased viral replication. Viral Immunol 2010; 23:557–65.

33. Ye J, Chen Z, Zhang B, Miao H, Zohaib A, Xu Q, et al. Heat shock protein 70 is associated with replicase complex of Japanese encephalitis virus and positively regulates viral genome replication. PLoS One 2013; 8:e75188.

34. Wells MF, Salick MR, Wiskow O, Ho DJ, Worringer KA, Ihry RJ, et al. Genetic Ablation of AXL Does Not Protect Human Neural Progenitor Cells and Cerebral Organoids from Zika Virus Infection. Cell Stem Cell 2016; 19:703–8.

35. Germi R, Crance JM, Garin D, Guimet J, Lortat-Jacob H, Ruigrok RW, et al. Heparan sulfate-mediated binding of infectious dengue virus type 2 and yellow fever virus. Virology 2002; 292:162–8.

36. Kurucz I, Tombor B, Prechl J, Erdo F, Hegedus E, Nagy Z, et al. Ultrastructural localization of Hsp-72 examined with a new polyclonal antibody raised against the truncated variable domain of the heat shock protein. Cell Stress Chaperones 1999; 4:139–52.

37. Lin RJ, Liao CL, Lin E, Lin YL. Blocking of the alpha interferon-induced Jak-Stat signaling pathway by Japanese encephalitis virus infection. J Virol 2004; 78:9285–94.

38. Khachatoorian R, Riahi R, Ganapathy E, Shao H, Wheatley NM, Sundberg C, et al. Allosteric heat shock protein 70 inhibitors block hepatitis C virus assembly. Int J Antimicrob Agents 2016; 47:289–96.

39. Howe MK, Speer BL, Hughes PF, Loiselle DR, Vasudevan S, Haystead TA. An inducible heat shock protein 70 small molecule inhibitor demonstrates anti-dengue virus activity, validating Hsp70 as a host antiviral target. Antiviral Res 2016; 130:81–92.

40. Taguwa S, Maringer K, Li X, Bernal-Rubio D, Rauch JN, Gestwicki JE, et al. Defining Hsp70 Subnetworks in Dengue Virus Replication Reveals Key Vulnerability in Flavivirus Infection. Cell 2015; 163:1108–23.

41. Crunkhorn S. HSP70 inhibitor blocks virus replication. Nature Reviews Drug Driscovery 2016; 15:18

